# Thermal asymmetries influence effects of warming on stage and size-dependent predator-prey interactions

**DOI:** 10.1101/2021.12.03.470870

**Authors:** Adam Pepi, Tracie Hayes, Kelsey Lyberger

## Abstract

Climate warming directly influences the developmental and feeding rates of organisms. Changes in these rates are likely to have consequences for species interactions, particularly for organisms affected by stage- or size-dependent predation. However, because of differences in species-specific responses to warming, predicting the impact of warming on predator and prey densities can be difficult. We present a general model of stage-dependent predation with temperature-dependent vital rates to explore the effects of warming when predator and prey have different thermal optima. We found that warming generally favored the interactor with the higher thermal optimum. Part of this effect occurred due to the stage-dependent nature of the interaction, and part due to thermal asymmetries. Furthermore, large differences in thermal optima between predators and prey (i.e., a high degree of asymmetry) led to a weaker interaction. Interestingly, below the predator and prey thermal optima, warming caused prey densities to decline, even as increasing temperature improved prey performance. We also parameterize our model using values from a well-studied system, *Arctia virginalis* and *Formica lasioides*, in which the predator has a warmer optimum. Overall, our results provide a general framework for understanding stage- and temperature-dependent predator-prey interactions, and illustrate that the thermal niche of both predator and prey are important to consider when predicting the effects of climate warming.

## Introduction

Predator-prey interactions are a key determinant of population and community dynamics and are expected to be affected by warming climate (Dell et al. 2014; Ohlund et al. 2014; Uszko et al. 2017). Climate warming has a broad range of effects on organisms, and many of these effects are due to the influence of temperature on metabolic rates in ectotherms (Gillooly et al. 2001; Brown et al. 2004; Dell et al. 2011). Increasing temperature, through increased metabolic rate, can increase movement speed, digestion rates, growth rates, and many other biological processes (Dell et al. 2011). The responses of these processes are commonly unimodal, increasing with higher temperature and then decreasing as biomolecular function starts to decline (Kingsolver 2009).

Warming can cause changes in equilibria and stability of predator-prey dynamics due to differences in temperature-dependence of predator and prey biological rates (Vasseur and McCann 2005; Uszko et al. 2017). Important changes to predator-prey dynamics may arise due to asymmetries in the thermal responses of predators and prey (Dell et al. 2014; Culler et al. 2015; Pepi et al. 2018; Davidson et al. 2021). A thermal asymmetry here refers to differences in thermal optima or thermal sensitivity that result in differential responses of predators and prey to changing temperature (Dell et al. 2014; Pepi et al. 2018). For example, when movement velocity responses to temperature are asymmetric in predators and prey, they can lead to nonlinear changes in the outcome of interactions that are dependent on foraging mode (Dell et al. 2014; Ohlund et al. 2014). An example of this includes predatory pike that are not fast enough to capture trout at low temperatures, but increase their attack speed with warming temperatures at a high enough rate that successful capture becomes possible above a certain temperature (Ohlund et al. 2014).

Temperature-dependent effects on predator-prey interactions can also result from asymmetries between prey growth rate and predator attack rates when predation is stage- or size-dependent. Lindmark et al. (2018) found that when predators are stage structured, warming can lead to a redistribution of biomass between life stages. Here, we explore when prey are stage structured. As the environment warms, prey growth rate speeds up, and the development time necessary to reach the invulnerable stage decreases, reducing the prey window of vulnerability to predators. At the same time, predators increase their attack rate with increasing temperature. The relative increase in prey growth rate vs. predator attack rate has the potential to determine whether net predation rate increases or decreases with temperature, constituting a potentially large indirect effect of changing temperature. For example, Ranchman’s tiger moth caterpillars (*Arctia virginalis*) are vulnerable to predation by ants when small, but become invulnerable when larger (Pepi et al. 2018) (Figure 1). With warming, increasing ant attack rate outpaced increased caterpillar development rate, resulting in a greater net predation rate (Pepi et al. 2018). Another example is that of Arctic mosquitoes, in which warming resulted in more rapid larval development and an increased attack rate by dytiscid predators, but in this case prey outpaced the predators, and warming resulted in increased prey survival (Culler et al. 2015). Similar results were found with mosquitos and dragonfly larvae in North Carolina, though the increased predation rate of one dragonfly predator was much less than another (Davidson et al. 2021).

**Figure 1.**
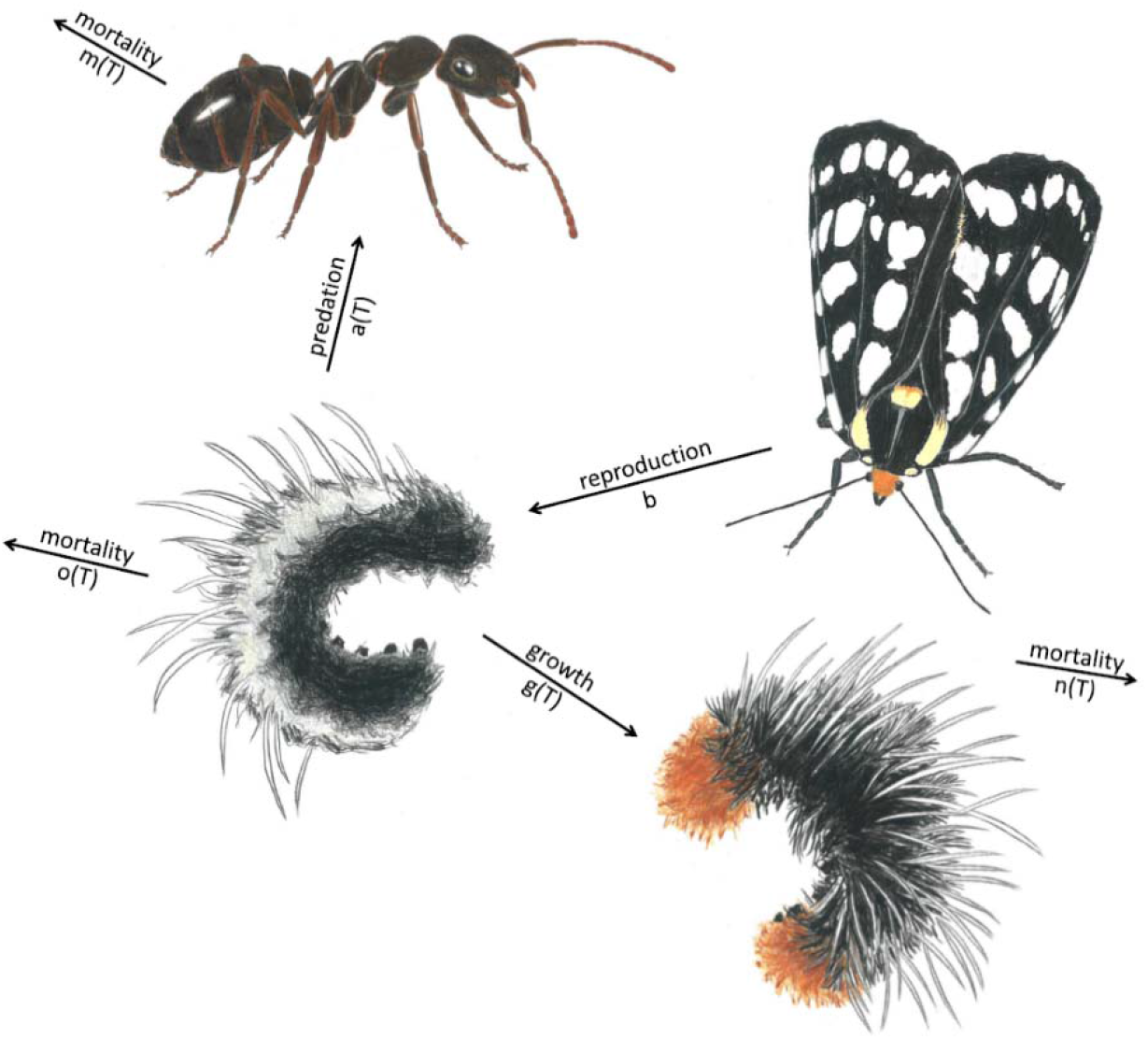
Conceptual figure illustrating temperature-dependent effects on a size-dependent predator-prey interaction. Small caterpillars (Juveniles, bottom left) are vulnerable to predation by ants (Predators, top left), but become invulnerable to ant predation when larger (bottom right) and when they become moths (top right). Here we model large caterpillars and moths as one group (Adults). Because predation and juvenile prey growth are temperature dependent, the effects of temperature depend on the ratio of their responses to warming.

Previous work has examined the consequences of thermal asymmetries in stage or size-dependent predation in a single specific system or using demographic models that represent only a single-generation (i.e., with no births or deaths; Culler et al. 2015; Pepi et al. 2018; Davidson et al. 2021). In the present work, we develop a multigenerational theoretical population model to examine the asymptotic effects of thermal asymmetries on stage or size-dependent predator-prey interactions. Our model allows us to make general predictions for the outcomes of these interactions with warming dependent on the thermal traits of interacting species. We further illustrate our model with an empirical parameterization using values from a well-studied system, *Arctia virginalis* (Lepidoptera: Arctiinae) and *Formica lasioides* (Hymenoptera: Formicidae).

## Methods

### Model description

To examine the effects of warming on a stage or size-dependent predator-prey system, we developed a population model of a prey species with vulnerable juveniles and invulnerable adults, and a predator species. We base the model on a simple Lotka-Volterra framework, with density dependence added to prey birth rates, and temperature-dependent growth, attack, and mortality rates. The resulting model is a Rosenzweig-MacArthur-type model (Rosenzweig and MacArthur 1963) with 2 prey stages and a Type I functional response, of the following structure:

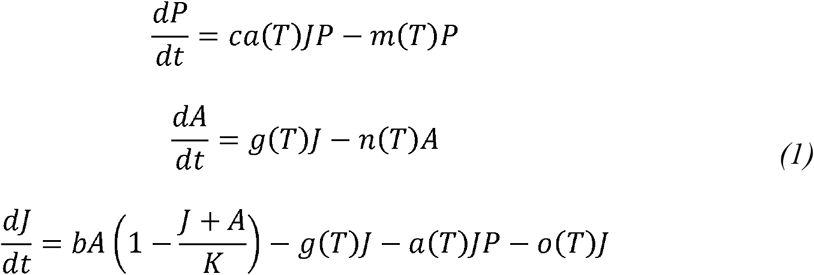

where *P* is the density of predators, *J* is the density of vulnerable juvenile prey, *A* is the density of invulnerable adult prey, *c* is the conversion rate of consumed juvenile prey to predators, *a*(*T*) is the temperature-dependent attack rate of predators on prey juveniles, *g*(*T*) is the temperature-dependent prey growth rate to adulthood, *b* is the birth rate of juveniles from adults, *K* is carrying capacity, and *m*(*T*), *n*(*T*), o(*T*) are the temperature-dependent mortality rates of predators, adult prey and juvenile prey respectively. We use a type I functional response for the sake of simplicity and tractability of the model. We leave *K* as temperature independent, as the temperature dependence of carrying capacity is the matter of some debate, and assumptions about whether *K* increases or decreases with temperature can have substantial effects on model findings (Uszko et al. 2017).

We analytically solved this model for equilibria. We also derived the Jacobian (eq. *S1*) which we used for numerical stability analyses (below).

### Temperature dependent parameters

For temperature-dependent attack rate and growth rate (*a, g*), we used a unimodal Gaussian function, from Taylor (1981):

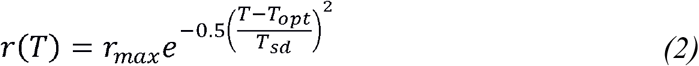

where *r*(*T*) is a temperature-dependent rate, *r*_*max*_ is the maximum rate, *T*_*opt*_ is the thermal optimum at which *r*_*max*_ occurs, and *T*_*sd*_ is the thermal niche breadth which controls the width of the function. Therefore, the function for each temperature-dependent parameter has three associated parameters: *r*_*max*_, *T*_*opt*_, and *T*_*sd*_ For mortality parameters (*m,n,o*)we used the following modified function:

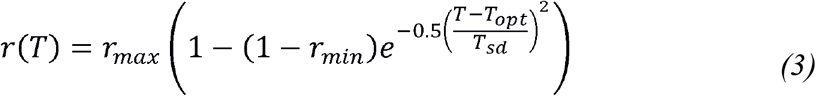

Such that *r*(*T*) was the lowest at *T*_*opt*_, increasing to a maximum mortality rate away from *T*_*opt*_. For mortality parameters, we varied *r*_*min*_ and kept *r*_*max*_ at 1 for the sake of simplicity (Table 1). *Thermal rate asymmetries*

**Table 1.**
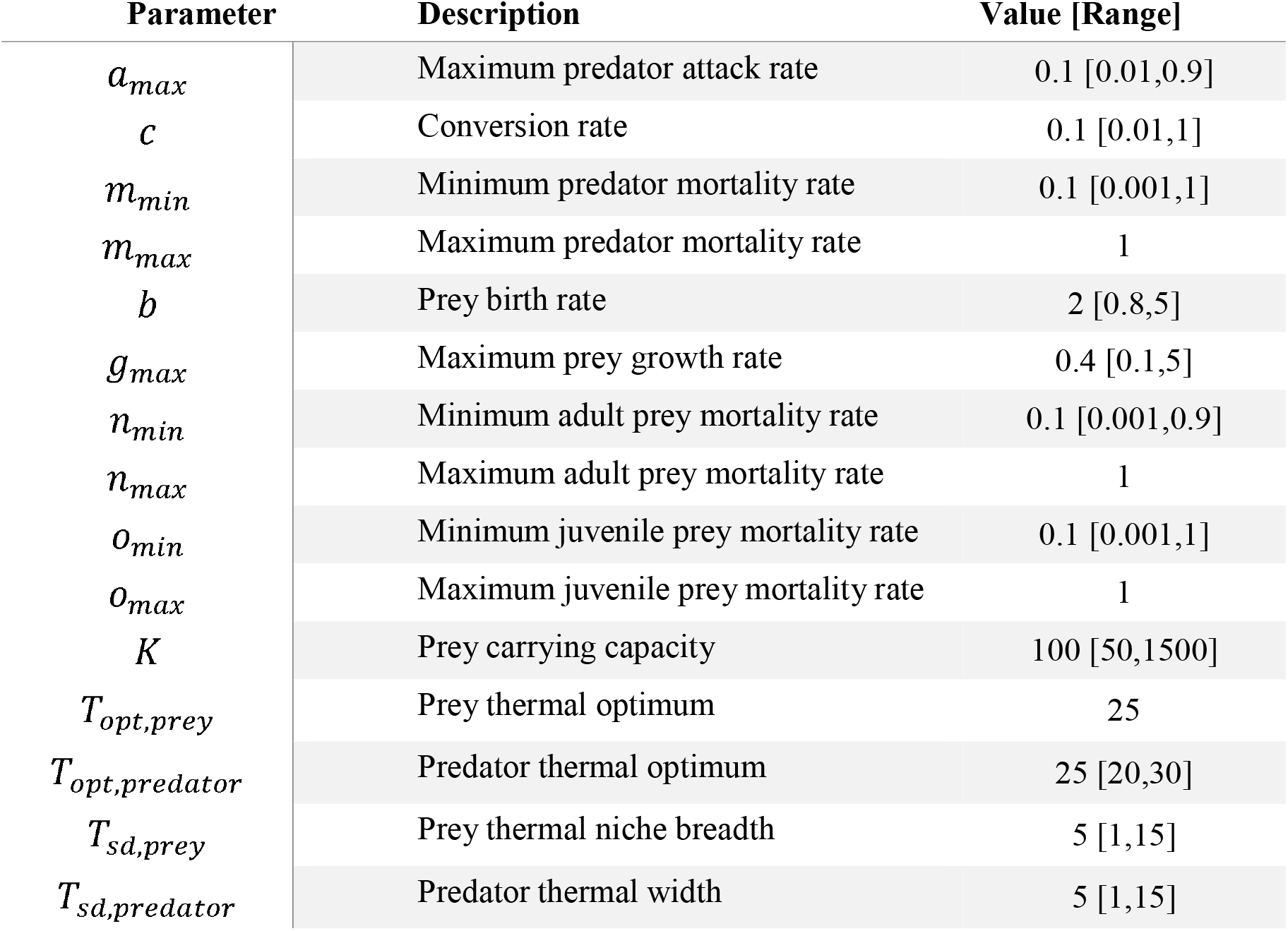
List of parameters in models, with description, value in base model, and range of values used in sensitivity analyses.

To examine the effects of asymmetries between predator and prey thermal rates, we conducted sensitivity analyses in which we numerically solved for equilibria using base model parameter values (Table 1), but varied predator thermal optima. We first present scenarios in which predator thermal optimum is the same as prey (both at 25°C), 2°C lower than prey, or 2°C higher than prey (Figure 2). We then examine the effect of a greater range of asymmetries, with predator thermal optimum ranging from 0-5°C greater than prey thermal optimum (Figure 3). For this second analysis, we present the change relative to the no-predator equilibrium, to separate indirect effects of temperature via the interaction with predators from effects solely due to the prey’s fundamental niche (i.e., prey temperature-dependent growth and survival). To estimate equilibria numerically, we used the function runsteady() from the package ROOTSOLVE (Soetaert and Herman 2008; Soetaert 2009), all using R (v. 4.0.1, R Development Core Team 2020).

**Figure 2.**
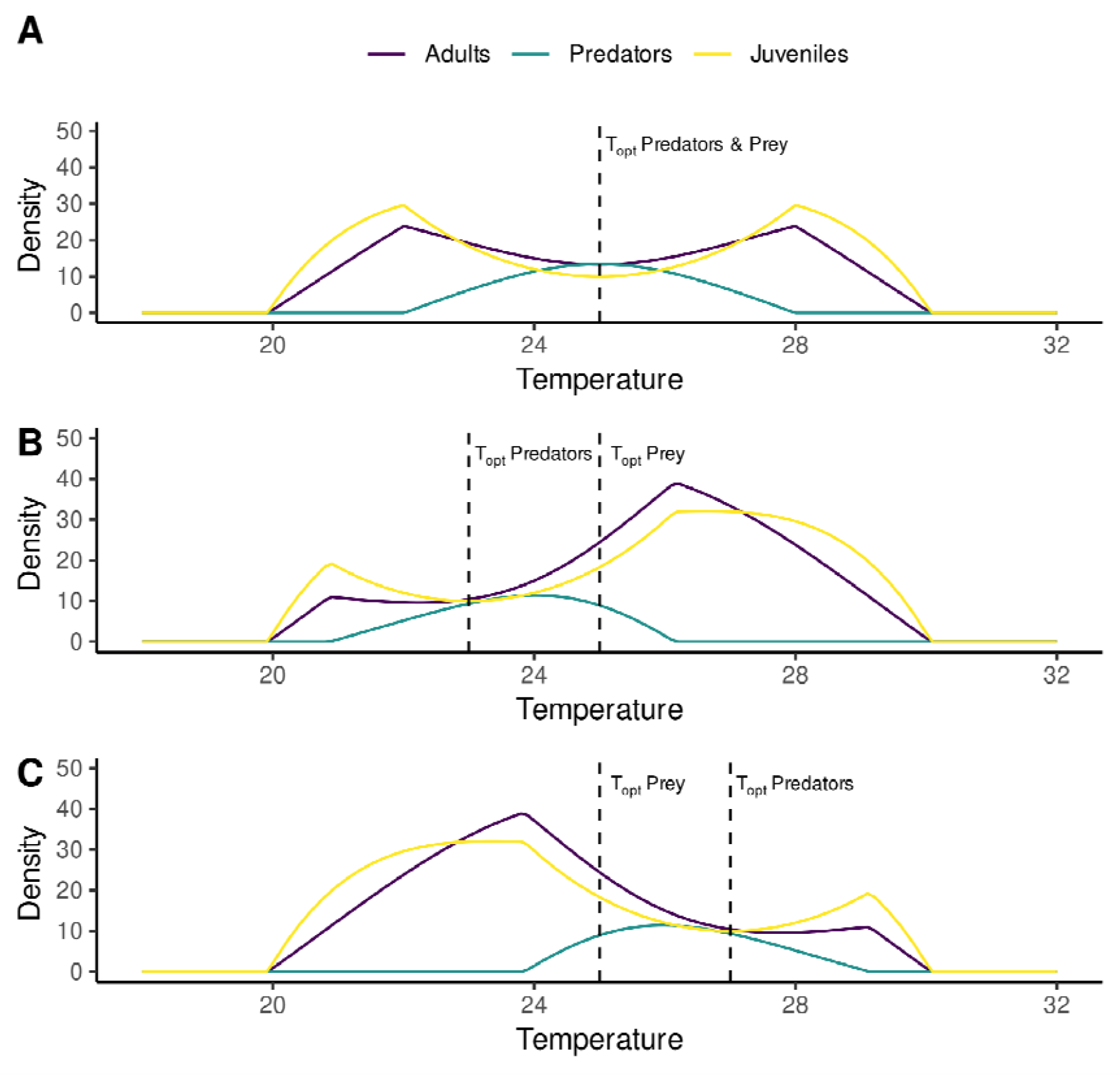
Model equilibria along a temperature gradient with base parameter values (Table 1), with no asymmetry between predator and prey thermal optima (A), predator thermal optimum 2°C below prey optimum (B), and predator thermal optimum 2°C above prey optimum (C).

**Figure 3.**
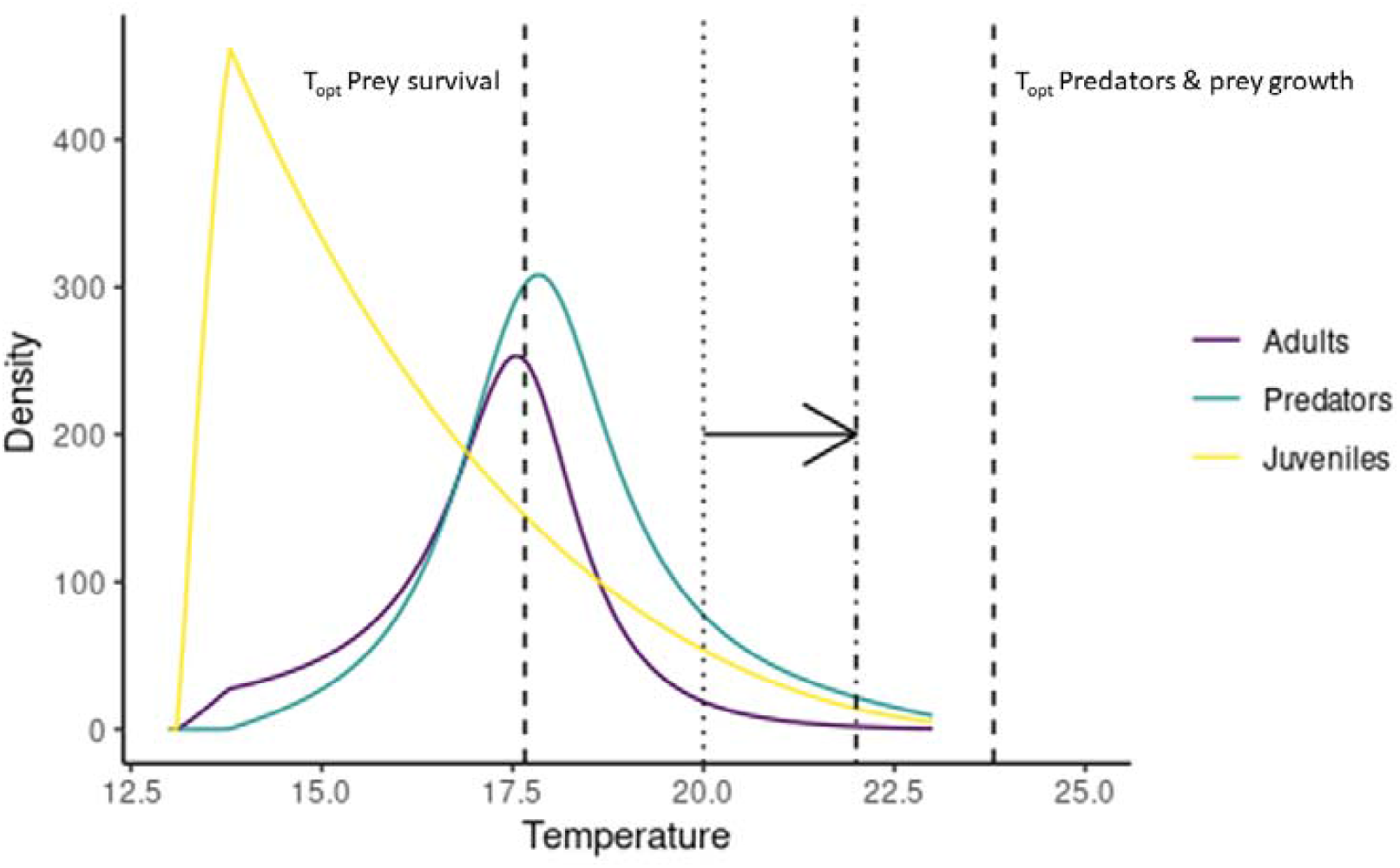
Model equilibria along a temperature gradient with parameter values from empirical data derived from *A. virginalis* (prey) and *Formica lasioides* (predator) (Table 2). Dashed lines indicate thermal optima of prey survival (17.66 °C) and both prey growth and predator attack rate (23.42 °C and 23.83 °C) Dotted and dot-dash lines and arrow illustrate a possible shift in temperature resulting from climate warming at the study site where these data were obtained, based on observed summer ground surface temperatures and 2°C of warming (Pepi et al. 2018). Over this range, densities are expected to decline, matching short-term predictions derived from previous experimental and simulation work (Pepi et al. 2018).

We also parameterized the model using empirical parameter estimates from the *Arctia virginalis* and *Formica lasioides* system (Table 2). We estimated prey temperature-dependent survival and growth from laboratory rearing data, and maximum predator attack rates based on field experiments (full methods in Pepi et al. 2018). We estimated predator thermal niche parameters based on field activity data from pitfall traps (full methods in Pepi and McMunn 2021). Minimum mortality rates for poorly studied stages of prey were based on the inverse of the normal length of that stage (based on field observations), and for predators, lifespan estimates for another species of the same genus from previous literature were used (Dussutour and Simpson 2012). Fecundity was also based on past estimates (English-Loeb et al. 1990). Detailed calculations are described in the supplement.

**Table 2.**
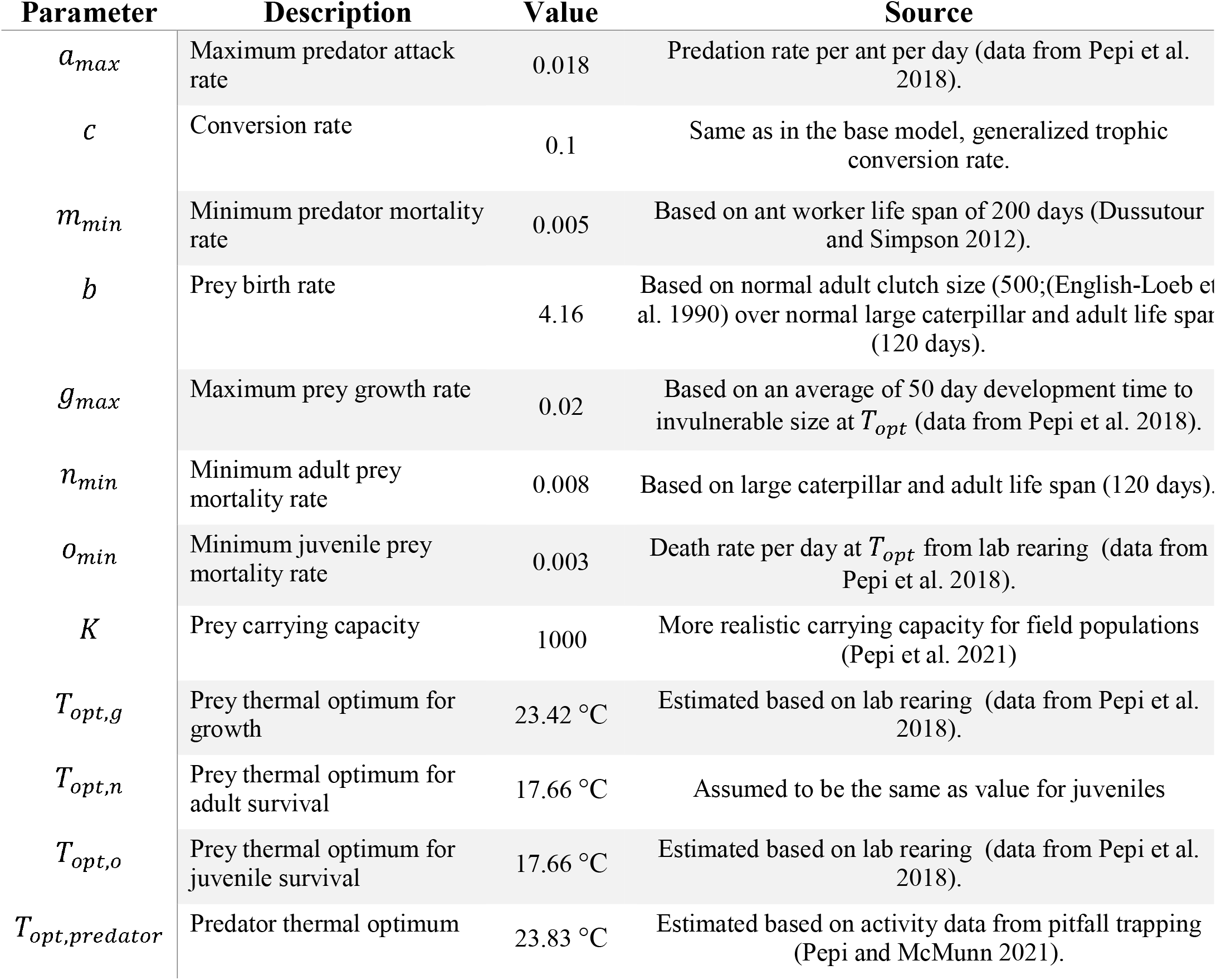
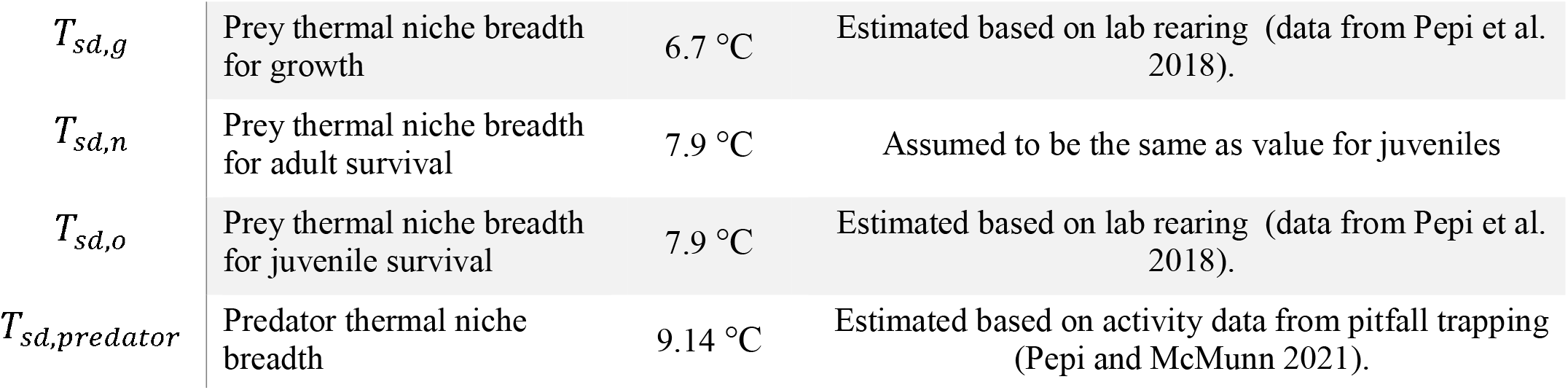
Empirical parameters values from *A. virginalis* (prey) and *Formica lasioides* (predator), with description, value, and source.

### Stability analysis

We conducted numerical stability analyses of the model as part of all analyses, because Routh-Hurwitz stability conditions derived analytically were sufficiently complex as to be uninterpretable (not shown). Using the Jacobian of the model (eq. *S1*), we extracted the leading eigenvalue with respect to each equilibrium resulting from the analyses of asymmetry and sensitivity analyses. We extracted leading eigenvalues using the R function eigen(), and considered all parameter combinations/equilibria with negative real parts to be stable even if imaginary parts were non-zero. Therefore, we considered both chaotic and stable limit cycle behavior to be unstable.

### Sensitivity analysis

We conducted single parameter sensitivity analyses of our model results. We conducted this in two ways: first, we conducted a simple sensitivity analysis in which we varied each parameter directly and plotted equilibrium prey and predator density for the range of values examined (Table 1, Figure S1). We also conducted a sensitivity analysis on the effect of asymmetry to test the generality of our main model result. We did this by looking at the change in equilibrium density of predators and prey having a thermal asymmetry of 2°C (predator *T*_*opt*_ = 27°C, prey *T*_*opt*_ 25°C), across a window of 2°C of warming. We used a 2°C window from 23-25°C, representing starting conditions below the thermal optimum. In general, we expect ectotherms to exist in thermal conditions mostly below their thermal optima under historical climate conditions (Deutsch et al. 2008).

## Results

### Model equilibrium

Analysis of the model resulted in the following equilibrium equations, for the equilibrium in which predators persist:

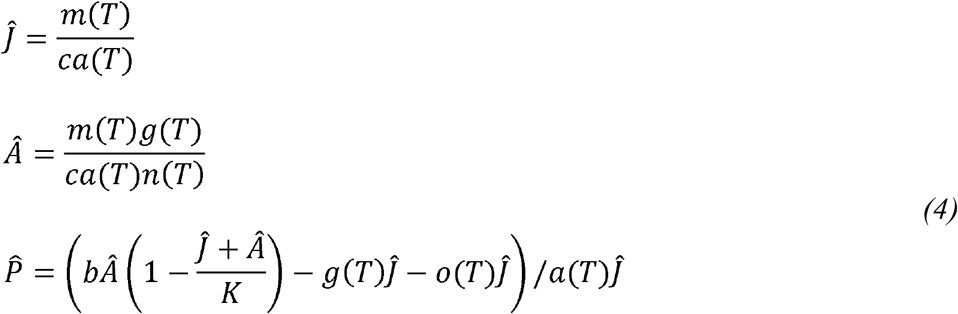

In this equilibrium, prey juvenile density depends on the ratio of predator mortality (*m*(*T*)) to attack rate (*a*(*T*)), and prey adult density depends on the ratio of prey growth (*g*(*T*))and predator mortality rate to predator attack and adult prey mortality rate (*n*(*T*)) The predator equilibrium is much more complex, and more easily interpreted through numerical results (Figures 2-5).

### Thermal rate asymmetries

In the base model with no asymmetry, prey equilibria were M-shaped with respect to temperature, with reduced densities in the region of predator persistence (Figure 2a). In the region of predator persistence and below the predator thermal optimum, warming caused prey densities to decline, even as the increase in temperature improved prey growth and survival rates. Thermal asymmetries resulted in directional (i.e., displaced from thermal optima) changes in prey equilibrium densities across temperature, so that there were overall increases or decreases with increasing temperature near prey thermal optima (Figure 2b,c). When predator thermal optimum was lower than prey thermal optimum, prey density increased with increasing temperature over an extended range of temperatures, including temperatures above the prey thermal optimum. The inverse was true when predator thermal optima were higher than prey thermal optima; prey density declined with increasing temperature over an extended range of temperatures, including temperatures below prey thermal optimum. Predator equilibrium densities increased with warming up to a temperature in between predator and prey thermal optima. Additionally, thermal asymmetries resulted in predator equilibria that were skewed in the direction of the predator thermal optimum.

In the parameterization of our model based on the *A. virginalis* and *F. lasioides* system, our results predicted that warming will cause a decline in juvenile, adult, and predator density. In this system, predators have a warmer thermal optimum than prey. However, the ambient temperatures historically occurring at the site (dashed line) were already above the prey thermal optimum before warming is simulated (dotted line), as data was taken from a location at the southern end of the prey’s range. Therefore, after warming, the inhospitable conditions for prey also drag down predator densities due to lack of food resources.

Next, we examined the effect of variable size of asymmetries by increasing the thermal optimum of the predator ranging from 0-5°C (Figure 4). With increasing asymmetry, the effect of predators decreased, as there is less overlap of predator and prey thermal niches. Reduced niche overlap thereby weakened the effect of predators on prey densities. Also, as expected, the negative impact of predators on prey is shifted to higher temperatures as asymmetry increases, resulting in a more directional (relative to prey thermal optima) but weaker impact of predators.

**Figure 4.**
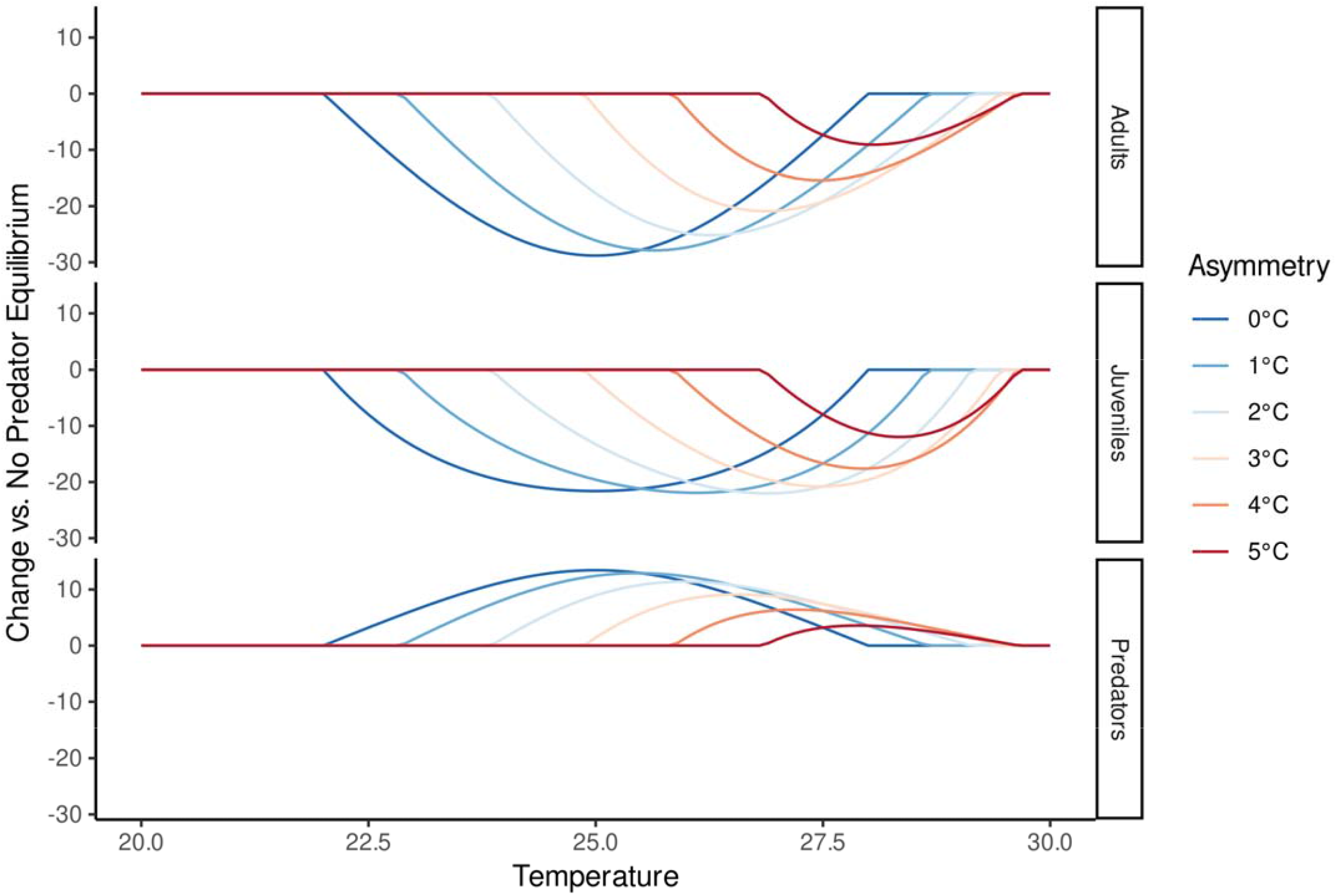
Illustration of effects of varying levels of asymmetry between predators and prey thermal optima on equilibrium densities of adults, juveniles, and predators (top to bottom) across temperature. Equilibrium densities are plotted as the change relative to the no-predator equilibrium, separating out temperature effects on prey due to thermal niche vs. changing interactions with predators. With higher degrees of asymmetry, predator effects become more strongly directional (i.e., displaced from prey thermal optima), but weaker as a narrower region of the predator and prey thermal niche overlap.

### Stability analysis

The coexistence equilibrium was stable across all parameter values analyzed (negative real part of leading eigenvalue), though in many regions predators did not persist (Figure 2, S1). Some regions exhibited damped oscillations, as signified by non-zero imaginary parts of eigenvalues (not shown).

### Sensitivity analysis

Sensitivity analyses show that the directional effects of asymmetric predator and prey thermal optima on equilibria persist across a wide range of parameter values (Figure 5). For many extreme parameter values, predators do not persist (gray regions, Figure S1). When predators do persist, there is a general pattern when predators have a 2°C warmer optimum than prey: warming leads to more predators (above the dashed line) and fewer prey (below the dashed line). The reduction in adult and juvenile prey densities is most extreme when attack rates are low, conversion rates are low, predator mortality rates are high, and carrying capacity is high. These larger reductions are mainly due to steeper local declines of prey across temperature because of a narrower predator niche (narrower range of persistence across temperature). The main result is generally disrupted if the maximum rates of many temperature-dependent parameters are extremely low or high, or *T*_*sd*_ values were too different from the base model. Changing *T*_*sd*_ disrupted detection of the main effect because varying the breadth of predator or prey thermal niches altered the M-shape or the position of declines in prey density such that prey declines did not always occur over the same interval of temperature (from 23-25°C).

**Figure 5.**
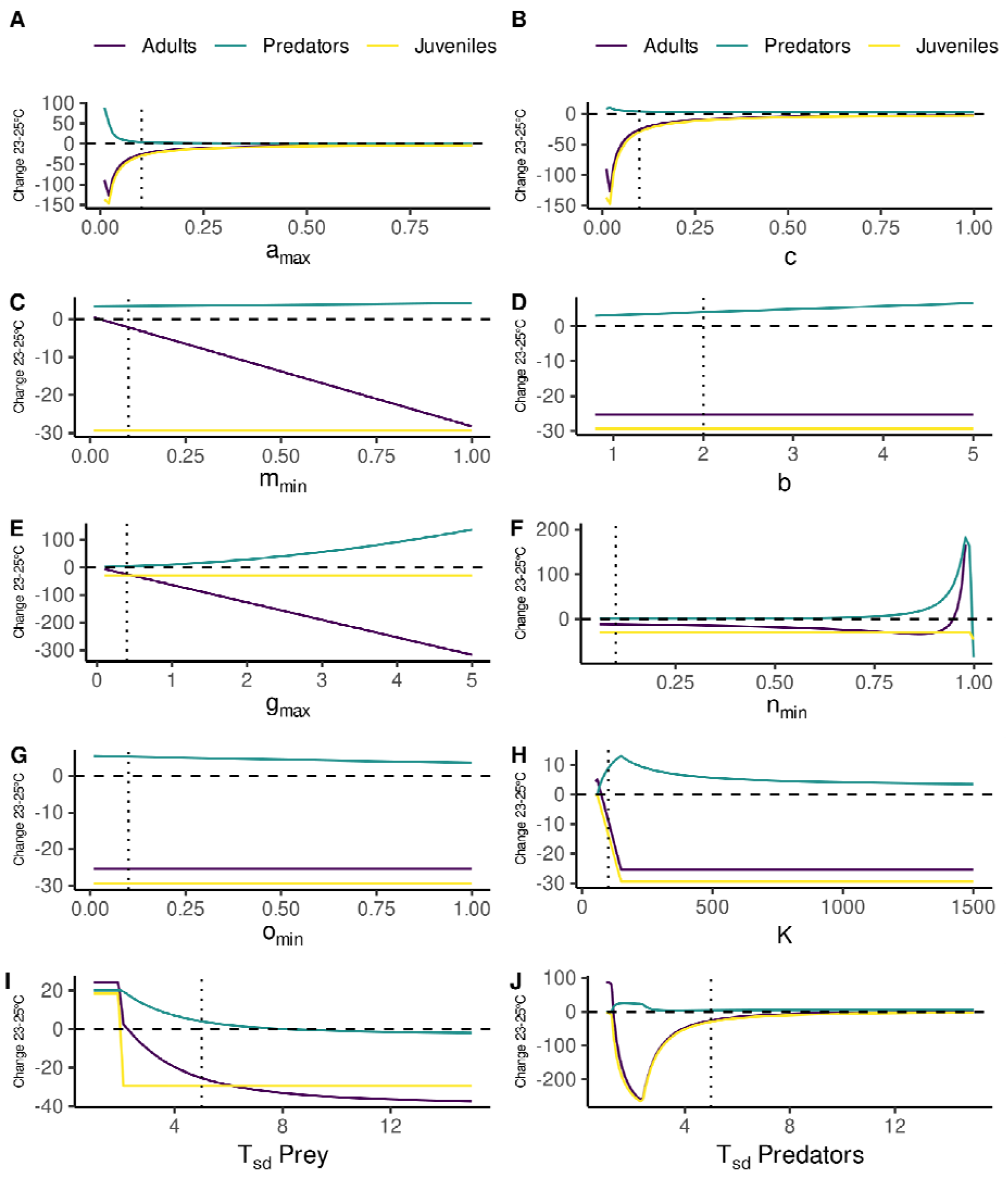
Single parameter sensitivity analysis of asymmetry effect on predators and prey with predator optimum 2°C higher than prey. Change in equilibrium density with 2°C of warming, from 23-25°C, is plotted. For panels C, F and G, which display mortality rates, one minus the minimum mortality rate is plotted on the x axis (i.e., survival: a value of one represents no mortality, and zero represents 100% mortality). Base parameter values are shown with dotted lines.

## Discussion

Overall, our model suggests that thermal traits of predators and prey, thermal optimum in particular, should predict the outcome of stage or size-dependent predator-prey interactions with warming. Specifically, if the predator thermal optimum is lower than prey thermal optima, we can expect prey populations to increase with warming (Figure 2B). Conversely, if predator thermal optimum is higher than prey thermal optima, we expect prey populations to decrease with warming (Figure 2C). Our modelling thus provides specific predictions for how we expect consumer impact on prey densities to change with warming depending on specific thermal traits of predators and prey. Previous work has found that prey survival may increase or decrease with warming (e.g., Culler et al. 2015; Pepi et al. 2018; Davidson et al. 2021) without providing specific insight into the factors which might cause a specific interaction to change one way or the other.

Interestingly, in the region of predator persistence below the predator thermal optimum, warming caused prey densities to decline, even as it brought temperatures closer to the prey thermal optimum. Conversely, in the region of predator persistence above the predator thermal optimum, warming led to increased prey densities, even as it brought temperatures away from the prey thermal optimum. This suggests that when predators persist, increases in predator attack rate with temperature outweigh increases in prey growth rate. Perhaps unsurprisingly, the effect of warming on predator densities depended on both the predator and prey thermal optima. Warming increased predator density up to a temperature between predator and prey thermal optima because predators thrive when both prey are abundant and attack rates are high. Our results provide clear support to a frequent call in recent ecological literature that interactions should be incorporated into projections of the ecological impacts of climate change and not solely the fundamental environmental niche (e.g., Araújo and Luoto 2007).

Our main findings were quite robust to a wide range of parameter values. However, some parameters had significant impacts on the main findings of the model. In particular, the degree of asymmetry between predator and prey thermal optima had the effect of decreasing consumer impact on prey densities, while also increasing the directional impact of warming on the interaction (Figure 4). That is, with an increasing asymmetry, predators consumed less prey at peak consumption, but the peak of consumption was more offset from the prey optimum. Changes in prey density due to predators were more uni-directional with warming (i.e., either increasing or decreasing with increasing temperature near the prey thermal optimum). The ratio of predator to prey thermal niche breadth (*T*_*sd*_) also affected the outcome: having predators with much broader thermal niches than prey reduced the main effect as there was less change in predation rate with temperature. Altering *a*_*max*_ or *c* affected overall consumer impact on prey density, and extreme values of minimum mortality rates or *g*_*max*_ also affected the outcome. Overall, the main finding that asymmetric thermal optima between predators and prey cause directional changes in equilibria with warming persisted across the majority of biologically plausible parameter values.

In developing our model, we took an approach of using the simplest possible formulation to allow the clearest possible interpretation of our main results. For example, we assume a type I predator functional response, whereas type II or type III is likely more realistic for many predator-prey systems. In addition, we assume a Gaussian response of temperature-dependent traits, whereas often temperature responses are skewed (e.g., Gibert and De Jong 2001). We also assume that the thermal optima of temperature-dependent vital rates are the same within each species when choosing parameter values for our main model, which is not the case in many species, including *A. virginalis*. Further modelling efforts could explore these additional levels of complexity and realism.

We chose to represent temperature dependence using Gaussian functions rather than Arrhenius-Boltzmann functions which have previously been used in modelling temperature dependence (e.g., Vasseur and McCann 2005), because we wanted a simple representation of a unimodal thermal response rather than commonly used monotonic functions. This assumption had important consequences for our results because the equilibrium density of prey depends in part on the ratio of predator and prey thermal responses (eq. 4; the ratio of *g*(*T*)/*a*(*T*)). The ratio of two offset Gaussian functions continually increases or decreases, which results in part of the directional impacts of warming on prey with warming in our results. By contrast, exponential functions will have constant ratios, resulting in no directional effects of warming on prey. Based on empirical evidence (e.g., in Pepi and McMunn 2021), the ratio of trait responses is likely to vary across temperature, and thus exponential functions will not appropriately represent the type of thermally-asymmetric interaction that we examine in this work.

For *A. virginalis* in particular, we can conclude that while warming will favor predators as they have a higher thermal optimum, warming will also weaken the interaction between the two species (Fig. 3). These predictions correspond with findings from experiments (Pepi et al. 2018) and time-series analyses of long-term population monitoring data (Pepi, Grof-Tisza, Holyoak and Karban, unpublished analyses) that warmer temperatures have negative impacts on *A. virginalis* in the drying California environment. Thus, in this system, our model provides a way to link mechanisms observed with short-term experimental data with longer-term trends, providing some evidence that predator-prey thermal asymmetries may be involved in climate change’s impacts on population dynamics.

Our model describes a mechanism for thermal asymmetries to impact predator-prey dynamics that is likely relevant to many species and interactions (although only part of our results are driven by stage-dependent effects). Stage- and size-dependent predation is common in a broad range of taxa, whether due to vulnerability of larvae to predation (insects and invertebrates; amphibians, Werner 1986) or gape-limited predation (common in many aquatic organisms; e.g., Paine 1976; Mittelbach 1981; Christensen 1996). Besides the previously highlighted examples (*A. virginalis* and *F. lasioides*; mosquitoe larvae and aquatic predators), our modelling approach is potentially relevant for many other well-studied predator-prey systems such as *Daphnia* and *Chaoborus* (Tseng and O’Connor 2015), salmonids (Ohlund et al. 2014), or grasshoppers and spiders (Laws and Joern 2013; Schmitz et al. 2016). Making accurate predictions will require a detailed understanding of the thermal responses of both predators and prey. Empiricists have documented the response of growth, mortality, and attack rates to increasing temperatures; however, we argue current knowledge is particularly lacking with respect to testing these vital rates at the upper end of species’ thermal limits.

Thermal asymmetries combined with ontogenetic shifts in vulnerability to predation may have important food web consequences beyond the predator-prey interactions studied here (de Roos 2021). There is the potential for more complex dynamics or alternative stable states to arise given that first, changes in temperature can have unpredictable and system-specific effects (Abbott et al. 2014); second, the presence of an invulnerable stage is known to influence stability and can lead to alternative stable states (Murdoch et al. 1987; de Roos and Persson 2002, 2013); and third, other trophic levels can affect the strength of predator-prey interactions. While not temperature-dependent, a previous model of stage-dependent predation with explicit resource dynamics found productivity reduced the strength of a predator’s impact (Chase 1999), a result parallel to our finding that a larger thermal asymmetry can reduce the strength of predator-prey interactions. Further work incorporating thermal performance of entire communities will offer further insight into the consequences of warming (e.g., Pepi and McMunn 2021).

## Supplement

### Jacobian

We found the Jacobian of the model, arranged as follows:

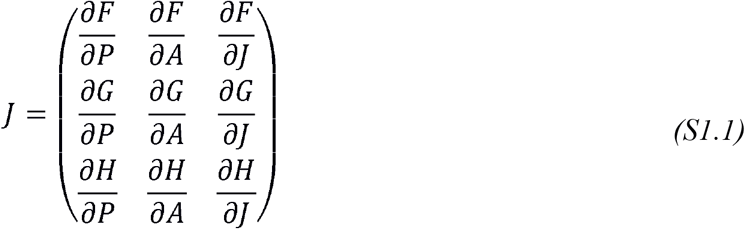

Where 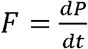 is the predator equation, 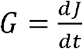 is the juvenile prey equation, and 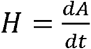 is the adult prey equation. After taking partial derivatives, the result is as follows:

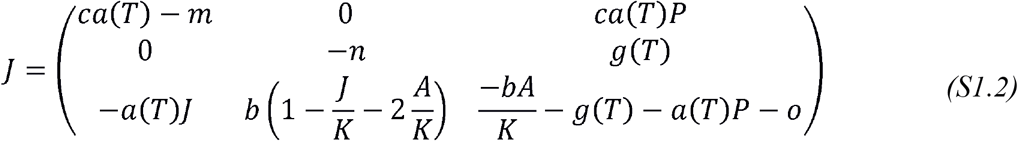

## Methods for parameterization

To estimate maximum attack rates, caterpillars eaten per day per ant predator per caterpillar was calculated based on field data from Pepi et al. 2018 collected at Bodega Marine Reserve. Using the model of caterpillar survival in the field, model-predicted means on the response scale were extracted using lsmeans (Lenth 2019). Mean predicted survival over the experiment was converted to mortality (1-survival), multiplied by the number of caterpillars placed (3) and divided by the number of days the experiment ran (3) and the mean ant abundance in tents (15.1).

To estimate temperature-dependent growth and survival rates, caterpillars were reared at temperatures from 15-30 C in 2017 and 2018 in incubators (Pepi et al. 2018). Caterpillars were weighed at the beginning, and periodically reweighed, and survival estimated. We used nonlinear least squares [nls() in nlme (Pinheiro et al. 2019)] to fit eq. 2 to growth and survival rates at different temperatures, giving us estimates of Topt, and Tsd for survival and growth, and max survival. Since growth rate should be in terms of days until the invulnerable stage is reached, we fit a gamma GLM of the days until a weight of 50 mg was reached (roughly the invulnerability threshold in Pepi et al. 2018), with an estimated mean time at the thermal optimum for growth of 48.5 days (giving a transition rate of 0.02/day).

To estimate the thermal niche of predators, we collected hourly pitfall samples along with hourly temperature data from a mechanical pitfall sampler in the Sierra Nevada (Pepi and McMunn 2021). We estimated Topt and Tsd for predators by fitting eq 2 to the temperature and activity data, using nonlinear least squares [nls() in nlme (Pinheiro et al. 2019)].

To calculate adult death rate, we used a maximum lifespan of predators of 200 days from Dussutour and Simpson (2012), which is roughly the maximum amount of time that workers of another species of Formica survived with optimum diet. To calculate birth rates, we used the mean number of eggs produced from the two years of data for unparasitized females from English-Loeb and Karban 1990. We based estimates of adult and large caterpillar lifespans based on observations and field notes, with large caterpillars visible in the field by March 1 most years, and adults being present into July in most years (∼120 days).

**Figure S1.**
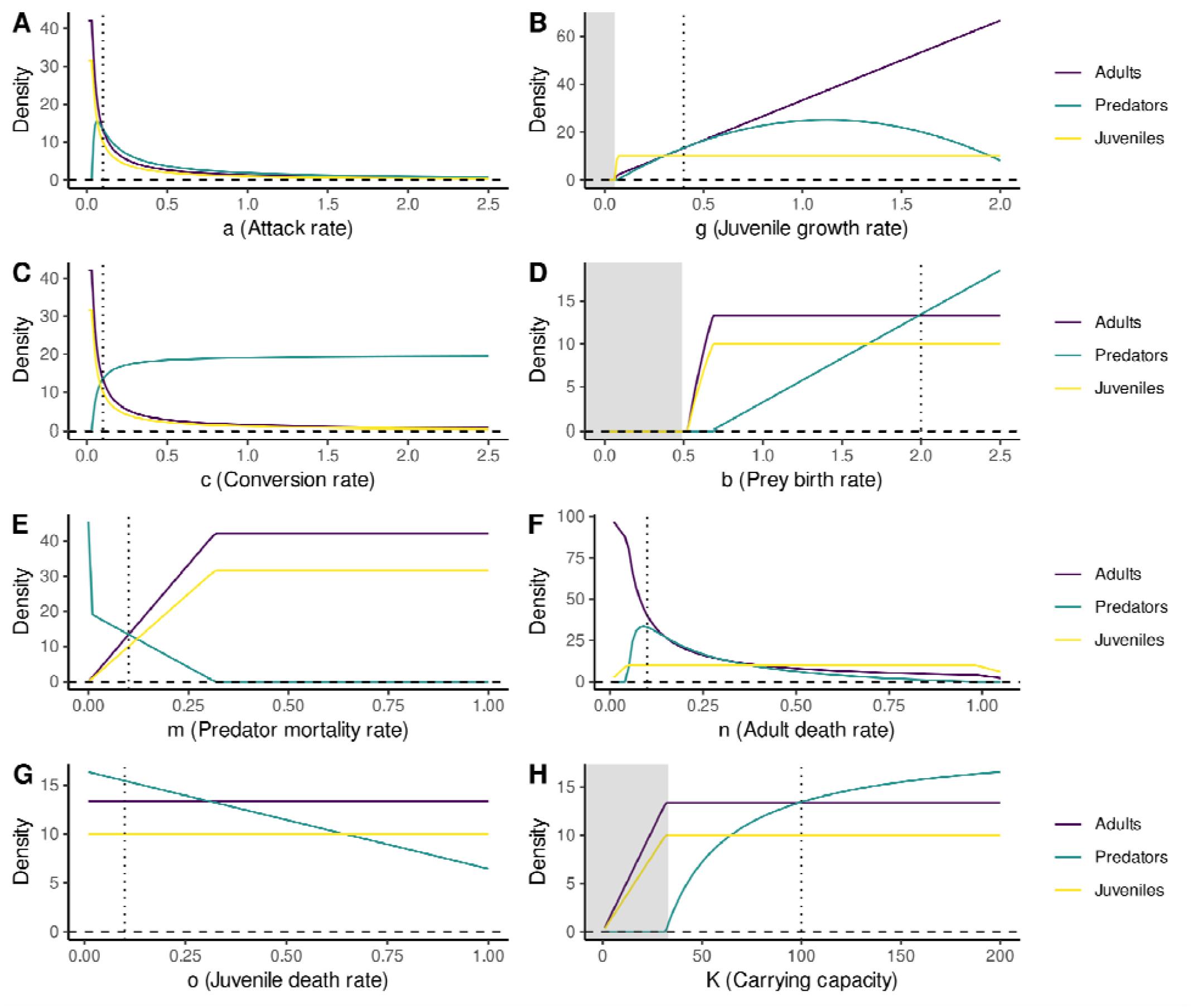
Single parameter sensitivity analysis, showing sensitivity of equilibria to parameters under the base model (Table 1). Regions where predators do not persist are shaded gray. Panels A, C and F are plotted with the y-axis on a log scale from improved visibility. Base parameter values are show with dotted lines.

## Notes

### Competing Interest Statement

The authors have declared no competing interest.

